# Combined flow cytometry and high throughput image analysis for the study of essential genes in *Caenorhabditis elegans*

**DOI:** 10.1101/218735

**Authors:** Blanca Hernando-Rodríguez, Annmary Paul Erinjeri, María Jesús Rodríguez-Palero, Val Millar, Sara González-Hernández, María Olmedo, Bettina Schulze, Ralf Baumeister, Manuel J. Muñoz, Peter Askjaer, Marta Artal-Sanz

**Author notes:** Equal contribution (Blanca Hernando-Rodríguez, Annmary Paul Erinjeri).

## Abstract

**Background:** The advancement in automated image based microscopy platforms coupled with high throughput liquid workflows has facilitated the design of large scale screens utilizing multicellular model organisms such as *Caenorhabditis elegans* to identify genetic interactions, therapeutic drugs or disease modifiers. However, the analysis of essential genes has lagged behind because lethal or sterile mutations pose a bottleneck for high throughput approaches.

**Results:** In *C. elegans,* non-conditional lethal mutations can be maintained in heterozygosis using chromosome balancers, commonly labelled with GFP in the pharynx. Moreover gene-expression is typically monitored by the use of fluorescent reporters marked with the same fluorophore. Therefore, the separation of the different populations of animals at early larval stages represents a challenge. Here, we develop a sorting strategy capable of selecting homozygous mutants carrying a GFP stress reporter from GFP-balanced animals at early larval stages. Because sorting is not completely error-free, we develop an automated high-throughput image-analysis protocol that identifies and discards animals carrying the chromosome balancer. We demonstrate the experimental usefulness of combining sorting of homozygous lethal mutants and automated image-analysis in a functional genomic RNAi screen for genes that genetically interact with mitochondrial prohibitin (PHB). Lack of PHB results in embryonic lethality, while, homozygous PHB deletion mutants develop into sterile adults due to maternal contribution and strongly induce the mitochondrial unfolded protein response (UPR^mt^). In a chromosome-wide RNAi screen for *C. elegans* genes having human orthologues, we uncover both, known and new PHB genetic interactors affecting the UPR^mt^ and growth.

**Conclusions:** A systematic way to analyse genetic interactions of essential genes in multicellular organisms is lacking. The method presented here allows the study of balanced lethal mutations in a high-throughput manner and can be easily adapted depending on the user’s requirements. Therefore, it will serve as a useful resource for the *C. elegans* community for probing new biological aspects of essential nematode genes as well as the generation of more comprehensive genetic networks.

## INTRODUCTION

Essential genes are critical for organismal development and are often associated with human diseases [1]. However, systematic analysis of essential gene function is being conducted at a slower pace than that of non-essential genes, in particular in multicellular model organisms as compared to yeast [2].

Approximately 30% of the genes in the *C. elegans* genome are essential [3, 4]. To investigate essential genes multiple approaches can be used that temporarily or partially reduce gene function [5]. Among them, temperature-sensitive (ts) alleles are extensively used, where protein function can be disturbed specifically upon temperature shift at any desired time. Although the number of identified ts alleles is rapidly increasing [6-9], not even 10% of the ~7,000 essential genes in *C. elegans* have yet a ts allele. An alternative to maintain and propagate lethal mutations is the use of balancer chromosomes. Around 85% of the *C. elegans* genome has been successfully balanced by large genomic rearrangements and community efforts aim at covering the complete genome [10-12]. Balancers prevent recombination with their homologous chromosomes and therefore, loss of lethal alleles in a population. Balancers carry alleles that negatively affect reproductive fitness when carried in homozygosis. In addition, an increasing number of chromosome balancers carry fluorescent transgenes, making the identification of homozygous mutant animals easy as they lack the fluorescent marker [11]. However, such strains are not easy to manage for large-scale analyses, as the isolation of the homozygous mutant population of interest is far too labour intensive.

One possible solution for the tedious task of selecting large numbers of homozygous worms could be automated worm sorting using the flow cytometer COPAS (Complex Object Parametric Analyser and Sorter) Biosort system (Union Biometrica, “worm sorter”) [13]. Recently, it has been utilized for sorting balanced mutants at the L3 [14] and L4 [15] larval stages to collect enough biological material for molecular biology techniques. Similarly, COPAS has contributed significantly to high throughput and high content analyses [16-18], like isolating homozygous young adults to perform image-based high-content assays to measure germ cell fate reprogramming [19]. The integration of automated worm sorting with microscopy platforms that facilitate automated image acquisition, worm segmentation and data analysis, has advanced existing genome-wide screening strategies [20-23]. Moreover, the small size and transparency of the worm coupled with the systemic RNAi methodology has facilitated high-throughput and high-content whole animal screenings using fluorescent protein reporters or dyes, as well as drug screening [24-29]. However, sorting of homozygous mutants from a balanced strain has never been used to study genetic interactions of essential genes.

Here we present an automated sorting and imaging strategy to screen for genetic interactions of essential genes whose loss-of-function alleles can be maintained using fluorescently-labelled chromosome balancers. In particular, we optimize a sorting protocol capable of distinguishing homozygous mutants that induce a GFP-stress reporter from balanced worms expressing GFP in their pharynx. Afterwards, we implement a segmentation protocol for image analysis that distinguishes eventual green pharynxes that can subsequently be discarded from the analysis using the Developer Toolbox software (GE, Healthcare). The protocol includes immediate background subtraction of all segmented worms and measures, in addition to fluorescence intensity, a variety of other parameters like area, length, curvature, etc. that can be defined by the user. We re-create the image analysis protocol using CellProfiler [30], a free and open source image-analysis software that caters for a variety of assays irrespective of the imaging system used. This protocol is easily adaptable to the user’s needs in terms of different fluorescent markers and image formats and resolutions. Doing so, we complement the features of the CellProfiler “WormToolbox”, a toolbox for high-throughput screening of image-based *C. elegans* phenotypes [23, 31], and make the protocol available to the broader community. We present validation that both protocols produce comparable results.

We provide proof of concept of the sorting and image analysis protocols by carrying out an RNAi screen in the mitochondrial prohibitin deletion mutant *phb-2(tm2998)*. The prohibitin (PHB) complex, a ring-like structure in the inner mitochondrial membrane, is composed of two subunits PHB-1 and PHB-2 [32]. Loss of either of the subunits leads to the absence of the whole complex, both in unicellular and multicellular eukaryotes [33, 34]. Prohibitins are strongly evolutionarily conserved proteins [35, 36], suggesting an important cellular function. While deletion of PHB does not cause any observable growth phenotype in the unicellular yeast *Saccharomyces cerevisiae* [37], in multicellular organisms, such as *C. elegans* [33] and mice [38], where it is ubiquitously expressed, the PHB complex is required for embryonic development. Postembryonic depletion of the complex by RNAi in *C. elegans* causes severe germline defects [33]. Despite the fact that their exact molecular function is yet to be deciphered [35, 36], they have been implicated in several age-related diseases [39, 40] and are involved in mitochondrial morphogenesis and maintenance of mitochondrial membranes by acting as scaffolds [41] or as chaperones that assist with protein folding and degradation [42]. Recently, PHB-2 has been described as being essential for Parkin mediated mitophagy [43]. Notably, homozygous prohibitin deletion mutants develop into sterile adults that strongly induce the mitochondrial unfolded protein response (UPR^mt^). Upon mitochondrial stress, cells respond by activating dedicated chaperones and proteases of the UPR^mt^ [44-46]. We use quantitative measurements of P*hsp-6::*GFP expression as a readout for the induction of the UPR^mt^, as well as worm size, with the aim of identifying genetic interactors of prohibitins and mechanisms modulating the UPR^mt^. We provide evidence that our methodology can detect both types of interactions.

Our protocol allows the characterisation of genetic interactions of essential genes, by applying high-content screening for a variety of phenotypes, including expression of a particular gene of interest (e.g. stress reporters). The combination of nematode sorting along with accurate high-throughput imaging further strengthens *C. elegans* as a powerful organism for the functional genomic analysis of essential genes.

## RESULTS

In the following sections, we demonstrate the usefulness of our sorting and image analysis protocol in identifying genetic interactions of essential genes, when mutants of these can be maintained using fluorescently labelled chromosome balancers. We first describe the phenotypes of mitochondrial prohibitin deletion mutants. We follow by describing in detail the sorting and imaging protocols used to perform systematic RNAi screens and providing experimental evidence for the wide applications of the protocol utilizing prohibitin mutants as example. Finally, we compare the performance of the Developer Toolbox protocol with the protocol based on the free and open source CellProfiler software.

### Characterisation of *C. elegans* mitochondrial prohibitin deletion mutants

In *C. elegans*, homozygous *phb-1* and *phb-2* deletion mutants produced by heterozygous mothers develop into adults due to maternal contribution but are sterile [47] and need to be maintained as balanced heterozygous strains. Here, the *phb-2(tm2998)* deletion is balanced using an inversion on chromosome II, *mIn1,* which carries an integrated pharyngeal GFP element, while for the *phb-1(tm2571)* deletion, we use a reciprocal translocation between chromosomes I and III*, hT2*, also accompanied by an integrated pharyngeal GFP element. Western blot analysis of *phb-1(tm2571)* and *phb-2(tm2998)* deletion mutants confirms the absence of the PHB complex (Figure 1a), as PHB-1 and PHB-2 are interdependent for protein complex formation and protein stability [33]. Therefore, *phb-1* and *phb-2* mutants show identical phenotypes. We use a method based on bioluminescence [48] to accurately measure the developmental timing of homozygous *phb-2* deletion mutants, which exhibit delayed development relative to wild type animals. All larval stages last approximately twice the time of wild type animals, with the third larval stage being more than two-fold increased. However, the duration of the molts is either not affected or mildly increased (Figure 1b). When assayed for longevity, PHB deletion mutants recapitulate the RNAi phenotypes [47], living shorter than wild type worms (Figure 1c).

**Figure 1:**
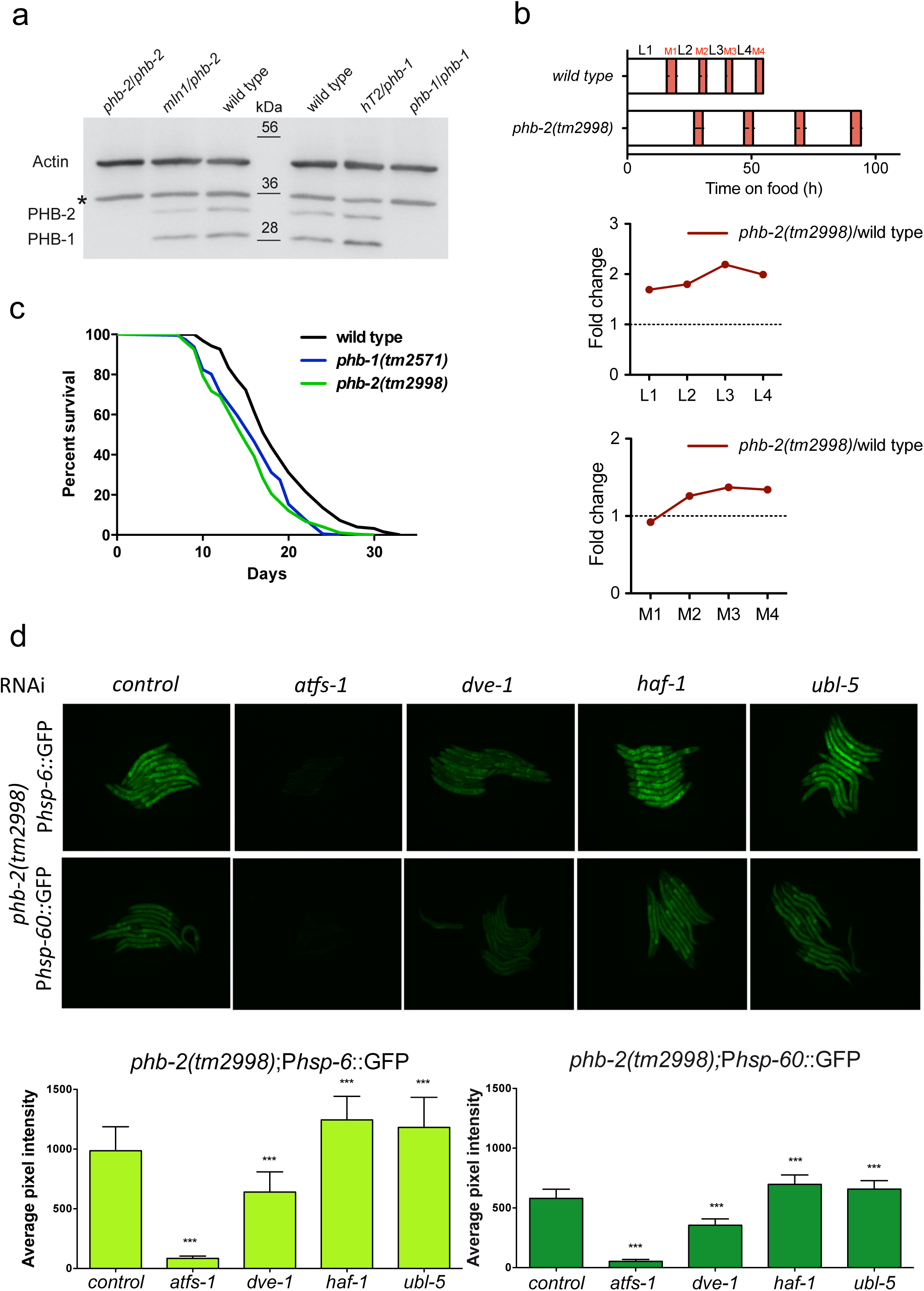
Characterization of *phb-2(tm2998)* mutants. **a.** Western blot showing PHB protein levels. Genotypes are indicated above the lanes. In homozygous *phb-1(tm2571)* and *phb-2(tm2998)* mutants both proteins, PHB-1 and PHB-2, are undetectable. Actin is used as loading control. Asterisk denotes unspecific band. **b.** Average duration of development for wild type (n = 12) and *phb-2*(*tm2998)* mutants (n = 18) at 20ºC, using the LUC::GFP bioluminescent reporter. Red represents the molts as inferred by low LUC signal. Error bars represent the SD of the duration of each interval. The graphs below represent the duration of larval stages (L1–L4, top) and molts (M1–M4, bottom) normalised to wild type. The duration of all larval stages (L1-L4) is significantly different between wild type and *phb-2(tm2998)* mutants (P value < 0.001; two-tailed unpaired t-test). The duration of the M2-M4 molting cycles are significant amongst wild type and *phb-2(tm2998)* mutants, with the exception of the M1 (P value < 0.001; two-tailed unpaired t-test). A representative experiment of two independent replicas is shown. **c.** Lifespans of wild type, *phb-1(tm2571)* and *phb-2(tm2998)* mutants. Lifespan of *phb-1(tm2571)* (mean = 17 ± 1 days, n = 215) and *phb-2(tm2998*) (mean = 14.5 ± 0.5 days, n = 302) are significantly shorter than the wild type (mean = 18 days, n = 137), P value < 0.0001, Log-rank (Mantel-Cox). Average of two independent assays is shown. **d.** Background dependent induction of UPR^mt^ reporters in prohibitin  deletion mutants. Fluorescent microscopy images of transgenic animals *phb-2(tm2998);*P*hsp-6*::GFP and *phb-2(tm2998);*P*hsp-60*::GFP treated with RNAi against the transcription factors ATFS-1 and DVE-1, the mitochondrial transporter HAF-1 and the ubiquitin-like protein UBL-5. Graphical representation of quantification of P*hsp-6*::GFP (bottom panel, left) and P*hsp-60*::GFP (bottom panel, right). The induced UPR^mt^ in prohibitin deletion mutants is suppressed upon depletion of *atfs-1* and *dve-1*, whereas the expression of both UPR^mt^ reporters is further increased upon depletion of *haf-1* and *ubl-5*. n > 30 in all conditions. **P value < 0.01, ***P value < 0.001, ANOVA test.

### PHB deletion induces the mitochondrial unfolded protein response by a non-canonical mechanism

Mitochondrial stress triggers the expression of the conserved mitochondrial chaperones HSP-6 and HSP-60, which have been used to screen for components of the UPR^mt^ signal transduction pathway [44-46, 49]. Nuclear genes encoding UPR^mt^ components required for HSP-6 and HSP-60 expression include the putative mitochondrial inner membrane ATP-binding cassette (ABC) transporter protein HAF-1 that exports the peptides resulting from unfolded proteins cleavage to the cytosol [45, 49]. Those peptides trigger an unknown signalling cascade that results in the nuclear localization of the bZIP transcription factor ATFS-1 [49, 50]. Additional transcriptional regulators of the UPR^mt^ are the transcription factor DVE-1 [45] and the ubiquitin-like protein UBL-5, that physically interact [44].

RNAi depletion of either *phb-1* or *phb-2* strongly induces the UPR^mt^ [46, 51, 52]. To monitor the UPR^mt^ in PHB deletion mutants, we incorporated the UPR^mt^ reporters, P*hsp-6::*GFP and P*hsp-60*::GFP [46], in *phb-2(tm2998)* mutants. The transcription factors ATFS-1 and DVE-1 are required for full induction of the UPR^mt^ in *phb-2* mutants (Figure 1d). However, HAF-1 and UBL-5 are not necessary for the PHB-mediated activation of the UPR^mt^, instead, their depletion further increases the expression of both UPR^mt^ reporters (Figure 1d). We confirm this observation using *haf-1(ok705*) deletion mutants, where RNAi depletion of either *phb-1* or *phb-2* increases the UPR^mt^ significantly more than in otherwise wild type animals (Additional file 1: Figure S1a). Similar to *haf-1(RNAi)* treatment, *haf-1(ok705)* deletion enhances the P*hsp-6::*GFP expression in *phb-2(tm2998)* deletion mutants (Additional file 1: Figure S1b). This suggests that HAF-1 not only is dispensable for signalling the UPR^mt^ upon prohibitin depletion, but blocking peptide transport through HAF-1 increases mitochondrial stress when depleting either subunit of the PHB complex. Similarly, we observe enhanced expression of P*hsp-6::*GFP in *haf-1(ok705*) deletion mutants upon RNAi against the mitochondrial AAA-protease *spg-7* (Additional file 1: Figure S1c), which cooperates with the PHB complex in mitochondrial quality control [53]. This observation is contrary to previously published data [49], but is in agreement with a previous report showing that *haf-1* is not required for induction of the UPR^mt^ caused by RNAi knockdown of *phb-2* [52].

Together, these data indicate that an alternative mitochondria-to-nucleus signalling mechanism might exist. We therefore develop an automated method to screen for PHB genetic interactors and regulators of the UPR^mt^. Our method consists of combined automated worm sorting and high-content image analysis that can be adapted and applied to any strain carrying a fluorescently labelled balancer.

### Sorting homozygous prohibitin deletion mutants

We use COPAS to sort homozygous *phb-2(tm2998)* deletion mutants from a mixed population of *phb-2(tm2998)/mIn1* balanced animals carrying the UPR^mt^ stress reporter P*hsp-6*::GFP. We sort homozygous *phb-2(tm2998)* deletion mutants at the second larval (L2) stage, to allow RNAi treatment during early development. This is more challenging than what has been accomplished earlier as we attempt to sort relatively small homozygous *phb-2(tm2998)* worms expressing P*hsp*-6::GFP all along their body from the population carrying the balancer *mIn1* and hence, expressing a pharyngeal GFP element but not the reporter P*hsp-6*::GFP.

The COPAS instrument measures the optical density of the object (extinction), the size of the object, as well as fluorescence intensity in three channels: green, yellow and red (Figure 2 and Figure 3a). Based on these parameters, the user can define criteria for sorting and dispensing the population of interest into multiwell microtiter plates. In order to make the sorting more accurate, the standard COPAS system is implemented with the Profiler II software. Instead of making a single integrated measurement of a signal, the Profiler gives a list of successive point measurements along the object passing through the flow cell, and builds a fluorescence profile. Based on these measurements, it can detect fluorescence intensity peaks along the length of the object (Figure 3b).

**Figure 2:**
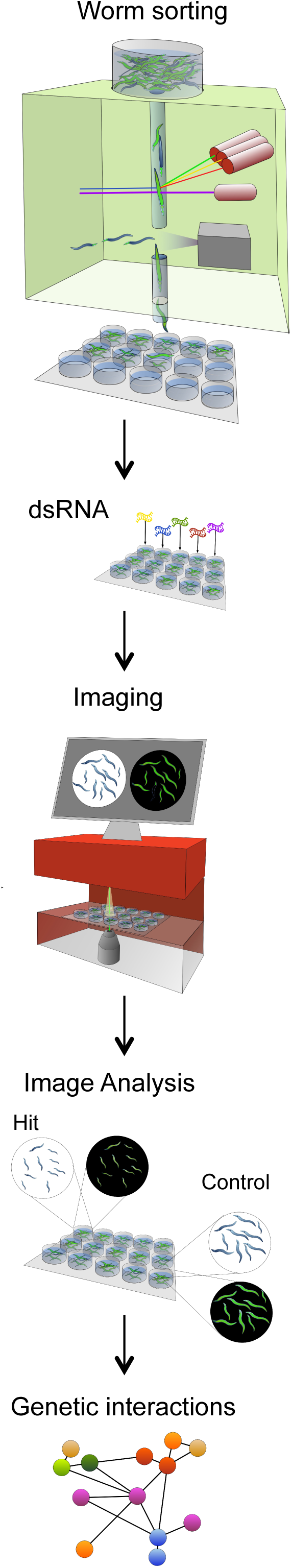
General strategy of the screen. Overview of the screening strategy for essential genes. Homozygous *phb-2(tm2998);*P*hsp6::*GFP mutants are sorted at L2 stage from a mixed population of balanced heterozygous *phb-2(tm2998)/mIn1* animals into multiwell plates using the COPAS Biosort, “worm sorter”. Next, bacteria containing the OrthoList RNAi sub-library are added to the wells and worms are incubated at 20ºC. When worms reach the desired stage they are imaged, in brightfield and fluorescent channels, using an automated microscope, IN Cell Analyzer (GE Healthcare). Using a user designed image segmentation protocol, hits can be defined based on different measurements like reporter expression or size of the worms in comparison with the control. Finally, hits can be analyzed by building genetic networks based on predicted and described interactions in different organisms.

**Figure 3:**
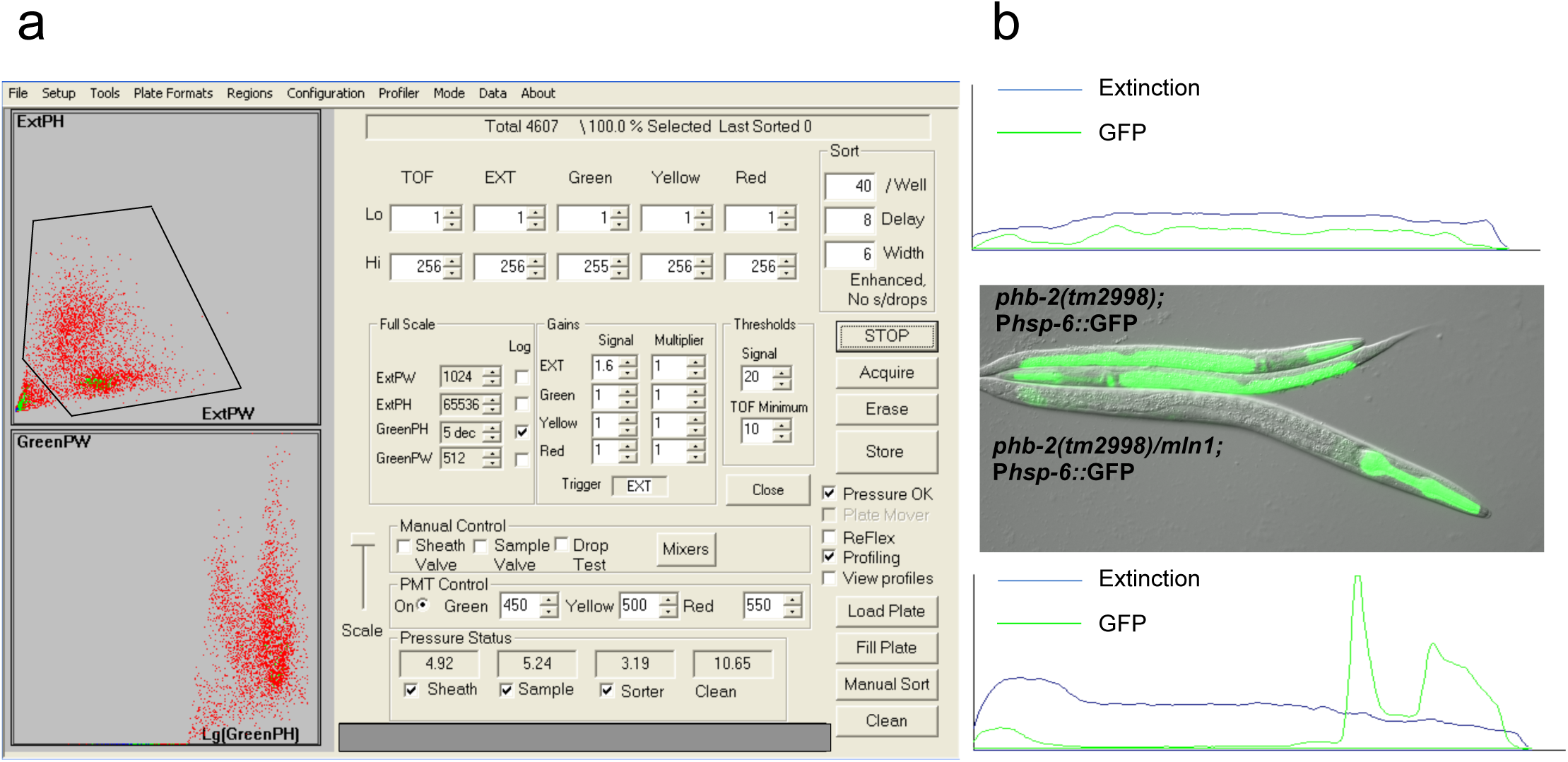
Worm sorting settings based on gating region and Profiler feature. **a.** COPAS Biosort conditions optimized for the sorting of homozygous *phb-2(tm2998);*P*hsp-6::*GFP at the L2 stage. The upper panel reflects the gating region based on the extinction peak height (ExtPH) and the extinction peak width (ExtPW) selecting the worm population distribution devoid of debris. The lower panel shows the worm distribution based on green parameters (green peak height (green PH) and green peak width (green PW). **b.** Utilising the Profiler II software to distinguish between *phb-2(tm2998)/mIn1;*P*hsp-6::*GFP and *phb-2(tm2998);*P*hsp-6::*GFP larvae using green parameters. The *phb-2(tm2998)/mIn1;*P*hsp-6::*GFP worms have a green peak height above 10,000 and are excluded (top panel). The *phb-2(tm2998);*P*hsp-6::*GFP larvae on the other hand having a green peak height ranging from 700-10000 and green peak width above 120 are accepted and sorted (bottom panel). In the picture we can see a balanced heterozygous *phb-2(tm2998)/mIn1;*P*hsp-6::*GFP expressing P*myo-2*::GFP in the pharynx and two *phb-2(tm2998);*P*hsp-6::*GFP homozygous deletion mutants with induced P*hsp-6::*GFP expression imaged at the moment of sorting, approximately 48 hours after synchronisation.

In our case, in order to remove small particles and possible debris and select specifically worms, a gating region is defined based on the extinction peak height (ExtPH) and the extinction peak width (ExtPW) (Figure 3a). Next, the green channel (500-520 nm wavelength) is used to select animals with green body from the rest of the population. As the green signal coming from the pharynx is more intense than the signal coming from the stress reporter, a signal range between 700-10,000 is accepted. These numbers refer to the highest value measured along the object. Nevertheless, in some cases, the signal coming from the pharynx is lower, thus a width criterion is added. To measure the width, the program integrates the widths of all the areas of the profile that exceed the value set up as the lower limit of peak height (700). Widths above 120 are accepted, excluding signals from the pharynx, which do not have the same extent as the body of the worm. All these numbers can be adjusted depending on the user requirements.

Once animals are sorted into 96-well plates, the bacteria producing double stranded RNA (dsRNA), prepared in parallel (see Additional file 2: Table S1 and Additional file 3) is added to the worms. In order to streamline the identification of relevant PHB interactors for human health, we assembled the OrthoList RNAi sub-library, maintaining the original name of the published compendium of *C. elegans* genes having human orthologues [54].

### Functional genomic analysis of chromosome I using the OrthoList RNAi library

*C. elegans* is an important invertebrate model for elucidating the mechanisms of conserved pathways relevant to human biology and disease. In *C. elegans*, RNAi can be applied by feeding worms bacteria expressing dsRNA for individual genes [56], which result in the generation of “feeding libraries” covering most of the predicted protein-coding genes in the genome [57, 58]. Of the ~20,000 predicted protein-coding genes in *C. elegans* [59], 7,663 genes have been listed in the “OrthoList” [54], a compendium of *C. elegans* genes sharing a human orthologue. Of those, about 80% (6,329 genes) are present in the Ahringer RNAi feeding library [57]. The generated “OrthoList RNAi sub-library” contains a total of 6,315 RNAi clones, since 14 clones from the Ahringer RNAi library did not grow during the preparation of the sub-library. The 6,315 bacterial clones are organized into 72 96-well plates, leaving empty the last column of the plates for the pertinent controls.

As a quality control for the OrthoList RNAi library, we sequenced 144 wide spread clones, of which 131 are identified correctly, corresponding to 91% of the bacterial clones being reliable. In 2011, Qu *et al.* performed an evaluation of the Ahringer *C. elegans* library [60], carrying out a bioinformatics analysis and resolved that 98.3% of the clones are trustworthy, even though 17.54% of the clones need to be re-annotated. Taking into account these numbers, we conclude that the OrthoList RNAi sub-library that we have generated is a high-quality tool.

Here, we present a functional genomic analysis of chromosome I using the OrthoList RNAi library. Chromosome I has 1,207 orthologous genes annotated, which corresponds to 14 96-well plates. RNAi is initiated at the second larval stage and animals are imaged two days later, at the young adult stage (Figure 1b).

### High-throughput image acquisition and analysis

After RNAi treatment, once worms reach the desired stage, animals are anesthetised and bacteria is washed off from microtiter plates before imaging (EL406 washer dispenser, BioTek). Whole-well brightfield and green fluorescence images are taken sequentially using the IN Cell Analyzer 2000 (GE Healthcare), an automated microscope designed for cell based high-content screening that we adapt for worm imaging using a 2x objective (Figure 4a). Image analysis is performed with a user-defined protocol within the Developer Toolbox software (version 1.9.2) (GE healthcare), accompanying the IN Cell Analyzer 2000 that enables direct upload and analysis of image stacks. The analysis protocol is built from individual target sets, which are linked together to obtain all the required measurement regions in one target. Each target is created stepwise: First, a pre-process is applied if required, for example to enhance image contrast prior to segmentation. Second, the target is segmented to generate a mask, and last, a sequence of post-processing operations is used to clean up the target mask. Once the targets are created and optimized, measurements for the individual targets or the linked target sets can be acquired on a per-worm basis such as size, shape or intensity in the different channels. In this protocol, we define the following targets; a well edge (used to subtract out any well debris), a worm, a dilated worm (used to calculate intensity of the background adjacent to the worm) and a worm with green head (so these can be segregated from the dataset downstream of image analysis). First, the well is segmented using the auto fluorescence of the well edge in the green channel, and after post-processing refinement steps, the resultant segmentation mask is inverted to give the well edge (Figure 4a - bottom). Next, the worms are segmented using the brightfield image; again, a pre-process is applied to create an enhanced image from which the worms are segmented (Figure 4a- top). After post-processing refinement of the worm mask, a subtraction of the well edge is carried out to remove well artefacts. Finally, in order to remove unwanted objects such as long fibres, air bubbles or overlapping worms, acceptance criteria based on morphological parameters are applied creating the worm mask (Figure 4a).

**Figure 4:**
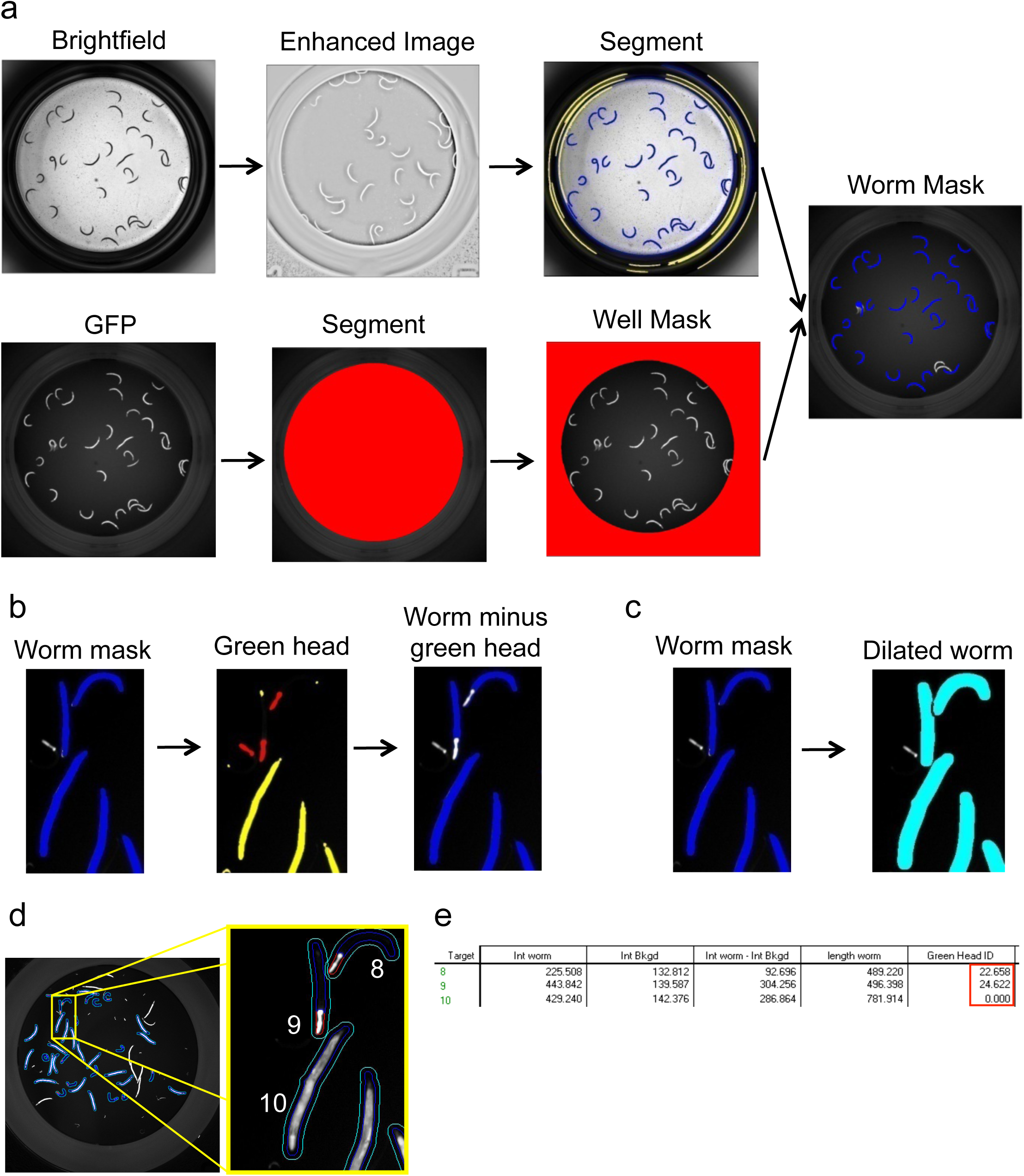
Outline of the GFP-image segmentation protocol for balanced mutants. **a.** Representative images acquired in brightfield and green channels. First, the well is segmented based on intensity in the green channel and then the image is inverted (bottom). The brightfield image is subjected to pre-processing to enhance the contrast for ease of segmentation. The segmented worms are tagged in blue and excluded objects are tagged in yellow (top). Exclusion criteria are specified in Additional file 4. The user can modulate these morphological parameters as per requirement. **b.** Identification of heterozygous animals (green head ID). Once worm mask is defined, green heads are segmented and tagged in red, excluded objects are tagged in yellow. Exclusion is based on area and intensity levels. The user can modulate these morphological parameters as per requirement. The next step subtracts the identified green heads from the worm bodies. As a result, segmented worms appear blue without the green heads, “worm minus green head”. **c.** Background subtraction. Segmented worms are dilated in order to facilitate measurement of the intensity of the immediate background. **d.** Target linking in order to compose a final target. Worm mask, dilated worm and worm minus green head targets are linked together. The final target shows the three measurement regions: the worms in blue, the immediate background in cyan and the green head in red. **e.** Identified targets with the corresponding measurements. Intensity of the worm (Int Worm), intensity of the immediate background (Int Bkgd), background subtraction (Int Worm - Int Bkgd), worm length and identification of green head (green head ID). Targets 8 and 9 correspond to heterozygous worms with a green head ID greater than 0. Additional measures like area, major axis length, X/Y position, form factor can be added as per user requirement during target linking.

Even though the sorting protocol is largely efficient, some heterozygous worms can be present in the well. Due to the longer developmental time of *phb-2(tm2998)* homozygous mutants (Figure 1b), the heterozygous worms lay progeny before the mutants reach the young adult stage, and several larvae with green pharynx can be found. The segmentation protocol is implemented by adding a step that identifies worms with green pharynx. Once worms are well delineated, green heads are segmented in the green channel image and acceptance criteria based in area and intensity are used to better define green heads (Figure 4b). Eventually, targets are linked to get all regions together. For two targets to be linked there must be at least one pixel overlap to achieve a linkage. The resultant is a target that has all three measurement regions: a worm mask, a dilated worm mask and a worm minus green head mask (Figure 4d). Worms with a green head are identified by the green head ID value (Additional File 4). A value greater than 0 indicates that a green head is present. A comprehensive list of measures, morphological and intensity based, for each worm is collected (Figure 4e). A break down of the protocol described here is provided as supplementary information (Additional file 4).

### Screen validation: PHB interactors affecting the UPR^mt^

In order to evaluate changes in GFP expression, an analysis pipeline consisting of several filtering steps, quality control and statistical test is computerized (see methods section). As shown in Figure 5a the negative (empty vector) and the positive (*atfs-1(RNAi)*) controls are clearly separable. Our RNAi screen is based on *C. elegans* genes sharing orthologues with humans annotated to Chromosome I. 208 genes are found to be necessary for the induction of the mitochondrial stress response while only the inactivation of one gene, *acd-1,* induces the reporter signal to a greater extent (Figure 5b). *acd-1* is an orthologue of members of the human SCNN (Sodium channels epithelial) family, which is involved in response to acidic pH and predicted to have sodium channel activity, based on protein domain information.

**Figure 5:**
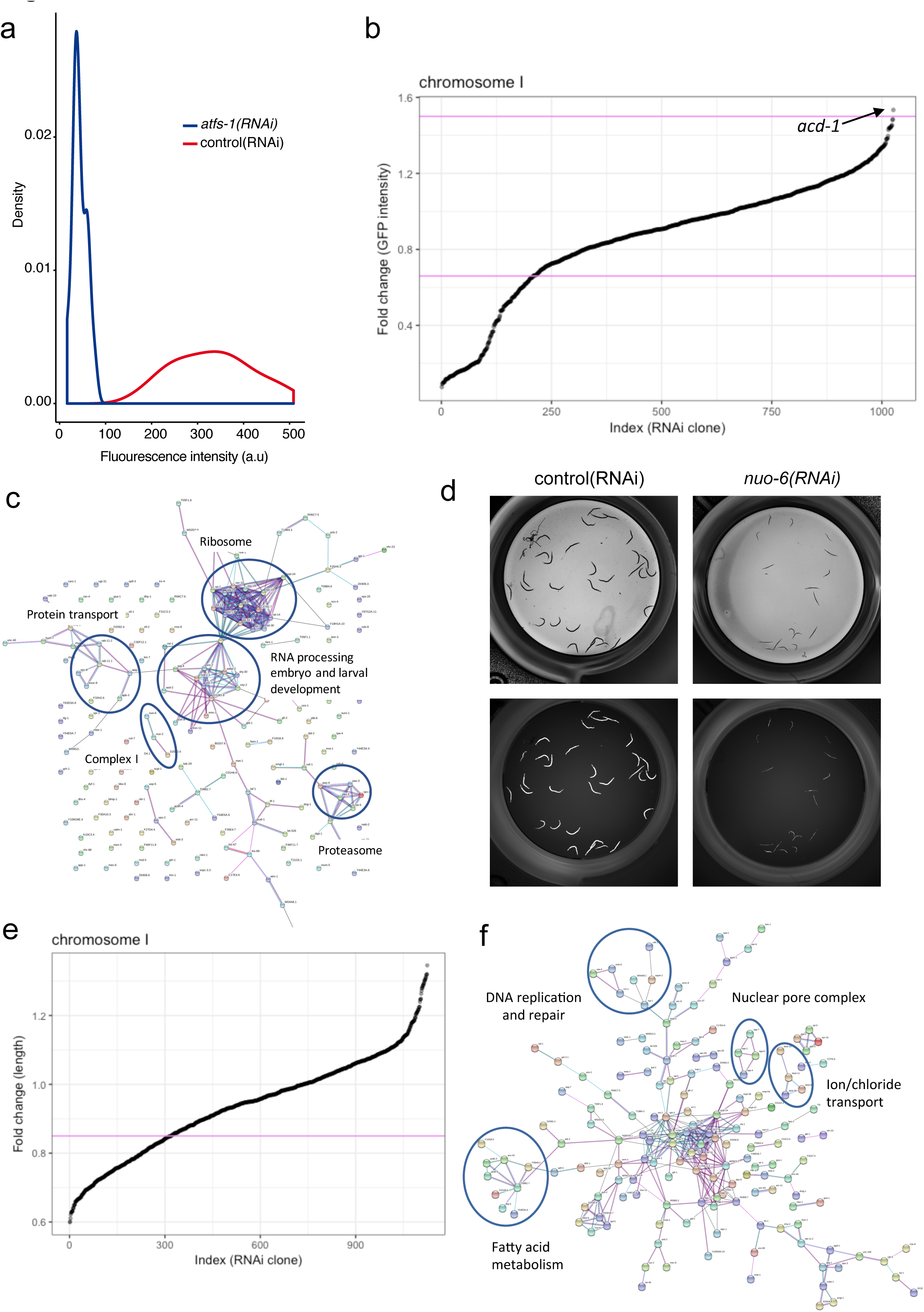
Analysis of the screening data. **a.** Frequency distribution of the negative controls (control *(RNAi)*) and positive controls (*atfs-1 (RNAi)*) based on the GFP intensities. **b.** Fold change (FC) of GFP intensity of the 1,207 tested RNAi clones against the ordered index. Genes with a P value < 0.001 and a FC < 0.66 or FC > 1.5 are considered as candidates. Depletion of 208 RNAi clones down regulates P*hsp-6*::GFP signal whereas only one candidate triggers a further induction of the reporter. **c.** Interaction network of the 208 genes which depletion reduces the UPR^mt^. Networks are built using String (http://string-db.org), based on predicted and described interactions in different organisms. Nodes are proteins and the edges represent the associations between nodes. Clusters show genes involved in processes previously described to be involved in the regulation of the mitochondrial stress response such as ribosome, proteasome, RNA processing, protein transport and complex I of the mitochondrial electron transport chain (ETC). **d.** Brigthfield and green images of control *(RNAi)* and *nuo-6 (RNAi).* Depletion of *nuo-6* triggers a developmental delay in addition to the reduction in the UPR^mt^ reporter expression. **e.** Fold change of worm length of the 1,207 tested RNAi clones against the ordered index. Genes with a FC < 0.85 are considered as clones affecting development. Depletion of 303 genes reduces size of the worms. **f.** Interaction network of the 303 genes which depletion reduces size of the worms. Networks are built using String (http://string-db.org), based in predicted and described interactions in different organisms. Nodes are proteins and the edges represent the associations between nodes. Clusters show genes involved in DNA replication and repair, fatty acid metabolism and ion channels and nuclear pore complex.

Under our conditions, we identify 45.5% of previously described genes required for the activation of the UPR^mt^ [44, 45, 49, 61, 62] (Additional file 5). Moreover, based on functional annotation clusters we find new genes never described to be involved in the regulation of the UPR^mt^ but belonging to pathways already described to suppress the mitochondrial stress response (Figure 5c and Table 1). We identify a large group of genes encoding proteins of the two ribosomal subunits, genes encoding proteins associated with protein transport including nuclear importins, genes encoding proteasomal subunits and genes involved in mRNA processing. In addition, we encounter a large number of genes involved in embryo and larval development. Interestingly, as shown in Figure 5c, we detect as well three genes encoding for subunits of mitochondrial NADH dehydrogenase complex (Complex I), *nuo-2*, *nuo-6* and *D2030.4*. Looking closely at these worms, we detect that they show a developmental defect (Figure 5d).

**Table 1:** Candidates reducing the P*hsp-6*::GFP signal and annotated to pathways previously described to be involved in the regulation of the UPR^mt^.

| Ribonucleoproteins |  |
| --- | --- |
| *rps_15* | 40S ribosomal protein S15 |
| *rps_19* | 40S ribosomal protein S19 |
| *rla_0* | 60S acidic ribosomal protein P0 |
| *C37A2.7* | 60S acidic ribosomal protein P2 |
| *rpl_1* | 60S ribosomal protein L10a |
| *rpl_13* | 60S ribosomal protein L13 |
| *rpl_17* | 60S ribosomal protein L17 |
| *rpl_24.1* | 60S ribosomal protein L24 |
| *rpl_7* | 60S ribosomal protein L7 |
| *mrpl_24* | Probable 39S ribosomal protein L24, mitochondrial |
| *snr_7* | Probable small nuclear ribonucleoprotein G |
| *snr_2* | Probable small nuclear ribonucleoprotein-associated protein B |
| *rpl_14* | Ribosomal protein, large subunit |
| *rpl_30* | Ribosomal protein, large subunit |
| *rps_10* | Ribosomal protein, small subunit |
| *rps_20* | Ribosomal protein, small subunit |
| Protein Transport |  |
| *apg_1* | AdaPtin, Gamma chain (clathrin associated complex 1) |
| *sec_8* | Exocyst complex component 4 |
| *ran_4* | Probable nuclear transport factor 2 |
| *bbs_9* | Protein pthb1 homolog |
| *rab_11.1* | Ras-related protein rab-11.1 |
| *nsf_1* | Vesicle-fusing ATPase |
| *apb_3* | hypothetical protein |
| *xpo_2/imb_5* | importin-beta-like protein |
| *imb_3* | importin-beta-like protein |
| Proteasome |  |
| *rpn_10* | 26S proteasome non-ATPase regulatory subunit 4 |
| *pas_3* | Proteasome subunit alpha type-4 |
| *pas_4* | Proteasome subunit alpha type-7 |
| *pbs_5* | Proteasome subunit beta type |
| *rpt_5* | proteasome regulatory partcile, ATPase-like |
| mRNA processing/splicing |  |
| *ncbp-2* | Nuclear cap-binding protein subunit 2 |
| *snr_7* | Probable small nuclear ribonucleoprotein G |
| *snr_2* | Probable small nuclear ribonucleoprotein associated protein B |
| *cel_1* | mRNA Capping Enzyme Like |

### Screen validation: PHB interactors affecting development

In addition to the quantification of P*hsp-6*::GFP expression, image analysis allows us to obtain many other measurements. In order to identify genetic interactors of PHB-2, we analyse the size of the worms in the same manner as described for the *hsp-6* reporter expression but filtering only based on the absence of green pharynx (green head ID = 0), keeping all worms irrespective of their size. In this case, we set up a threshold of FC < 0.85 to obtain the clones affecting size. We find 303 RNAi clones whose depletion leads to smaller size, probably due to developmental delays (Figure 5e). The Ahringer laboratory has performed multiple RNAi screens and assigned biological functions to many genes unknown until the present date [57, 63]. In addition, Ahringer and colleagues described many phenotypes, such as embryonic lethal or developmental delay, associated with RNAi depletion of individual genes in wild type worms. 31% of described RNAi clones showing a developmental delay in wild type animals also affect development of *phb-2* mutants (Additional file 6). We expect to have a developmental phenotype in *phb-2* mutants upon depletion of those genes that cause this phenotype in wild type worms, although it should be noted that the screening conditions are not the same. Fraser and Kamath [57, 63] subjected worms to RNAi from eggs, whereas we do it from the second larval stage. As expected, among the genes affecting development we find genes encoding ribosomal subunits, proteasome subunits and genes involved in protein transport. Moreover, we identify many genes implicated in fatty acid metabolism, some of them encoding for mitochondrial proteins such as *T08B2.7, lbp-5, F08A8.3, acs-16, acdh-3, acdh-4, F10G8.9* (Additional file 6). Interestingly, by doing protein-protein interaction analysis (Figure 5f) we describe a cluster of genes encoding for nuclear pore complex proteins, *npp-2, npp-4, npp-6, npp-7*, as well as a cluster of genes involved in DNA replication and repair, *cdt-1, aspm-1, icp-1, crn-1, msh-6* and *rpa-4*.

### Additional features of the imaging protocol: analysis in the red channel

Furthermore, we put forward an image acquisition and segmentation protocol that incorporates intensity measurement in the red channel into the above described image acquisition and segmentation protocol for balanced mutants (Figure 6). For this protocol, sequential images of whole wells in brightfield, green and red channels are acquired with the IN Cell Analyzer 2000 using the 2x objective. Image analysis is facilitated by a slightly modified version of the user-defined segmentation protocol described above for GFP images. This modified protocol follows the same initial steps of well segmentation in the green channel image and segmentation of worms in the brightfield image, followed by subtraction of the well edge, detection and identification of heterozygous worms with pharyngeal GFP and finally, dilation of the worm for measurement of the immediate background (Figure 4a-c and Figure 6a). Acceptance criteria based on morphological parameters are applied to remove unwanted objects. Target linking is done to achieve a final target set encompassing the final worm mask, the dilated worm and the worm minus green head linked together (Additional file 7: Figure S2). Defined measures are collected for each worm covering morphological and intensity based measurements from both the green and the red channels (Figure 6b). A break down of the protocol is provided in the Additional file 8.

**Figure 6:**
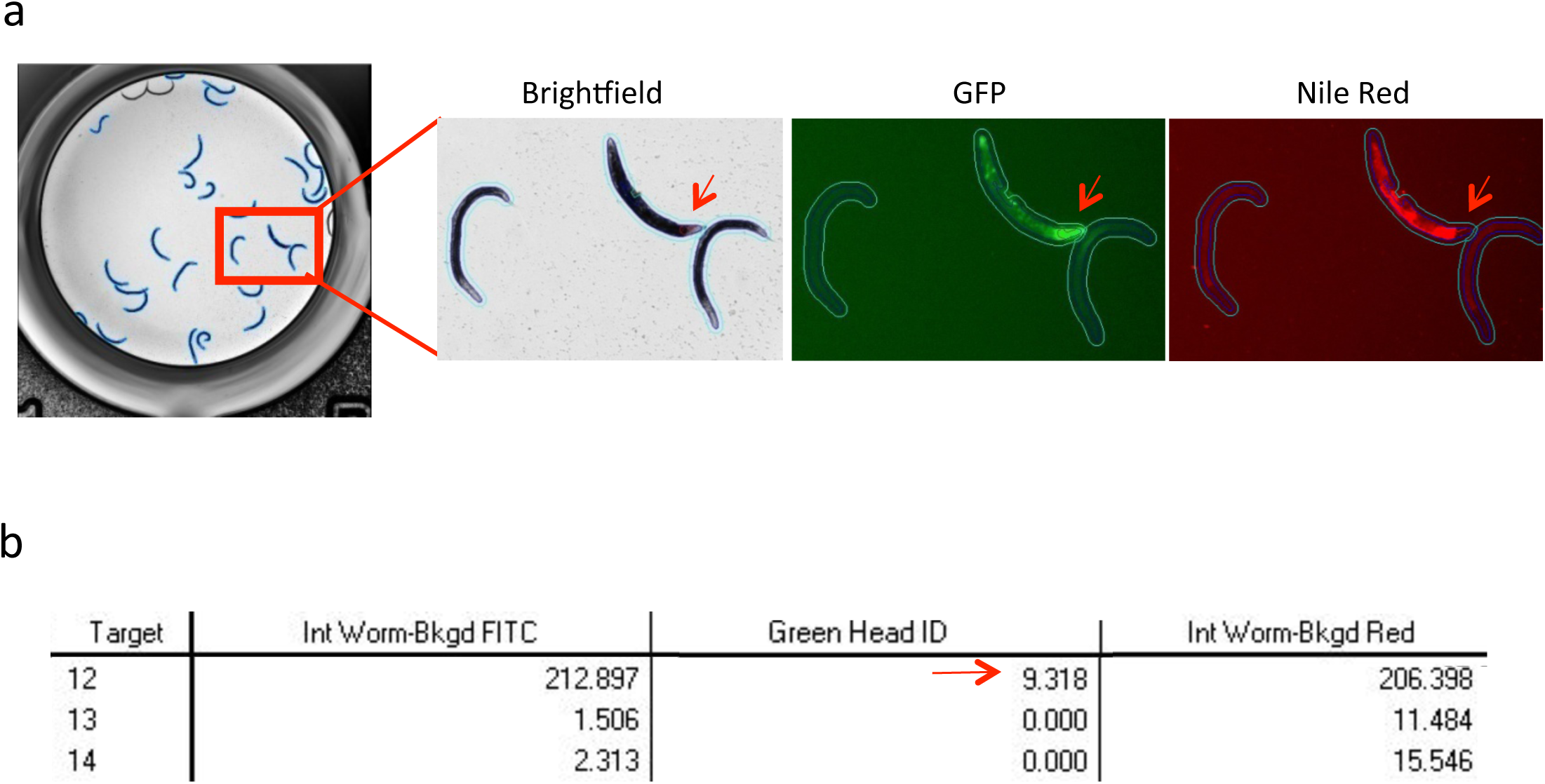
Combined green/red image analysis protocol. **a.** Images acquired in brightfield, green and red channels with the IN Cell Analyzer 2000 (GE Healthcare), 2x objective. Linked targets outline after worm, immediate background and green head detection (worm – blue, immediate background – cyan, green head – red). A balanced heterozygous animal expressing pharyngeal GFP has been identified (red arrow) within a population of homozygous *phb-2(tm2998)* mutants. The heterozygous animal exhibits increased Nile Red staining in comparison to the homozygous *phb-2(tm2998)* mutants. **b.** Segmentation analysis output of the green/red image analysis for balanced mutants. Intensity of the worm subtracting the background intensity in the green image (Int Worm - Bkgd FITC), identification of green head (green head ID) and intensity of the worm subtracting the background intensity in the red image (Int Worm - Bkgd Red) are depicted in the analysis output. Target 12, with a green head ID greater than 0, corresponds to the heterozygous worm. Additional measures like area, major axis length, X/Y position, form factor can be added as per user requirement during target linking.

### Open sourcing of the segmentation protocol through CellProfiler

The segmentation protocols presented above are only compatible with the Developer Toolbox software and hence, usage would be limited to researchers with access to the GE platform. To circumvent this, we implement the protocol in the free and open source CellProfiler software, providing an analysis pipeline identifying green head worms, and measuring intensities in the green and red channels.

We compare worm segmentation outputs resulting from the Developer Toolbox and CellProfiler and find them comparable (Figure 7a). Intensity measures obtained from the Developer Toolbox and CellProfiler in the green and red fluorescence channels are also compared. For the red channel, we compare wild type and *phb-2(tm2998)* mutants after Nile Red staining. Depletion of PHB by RNAi in wild type animals results in low Nile Red Staining [47]. Similarly, we observe lower Nile Red staining in *phb-2* mutants irrespective of the software used for segmentation (Figure 7b). For the green channel protocol, we randomly select one plate from the RNAi screen performed using *phb-2(tm2998);*P*hsp-6*::GFP animals and re-analyse the images using the CellProfiler software. After data processing, the candidates resulting from both analyses are highly similar (Figure 7c), thus, validating the method.

**Figure 7:**
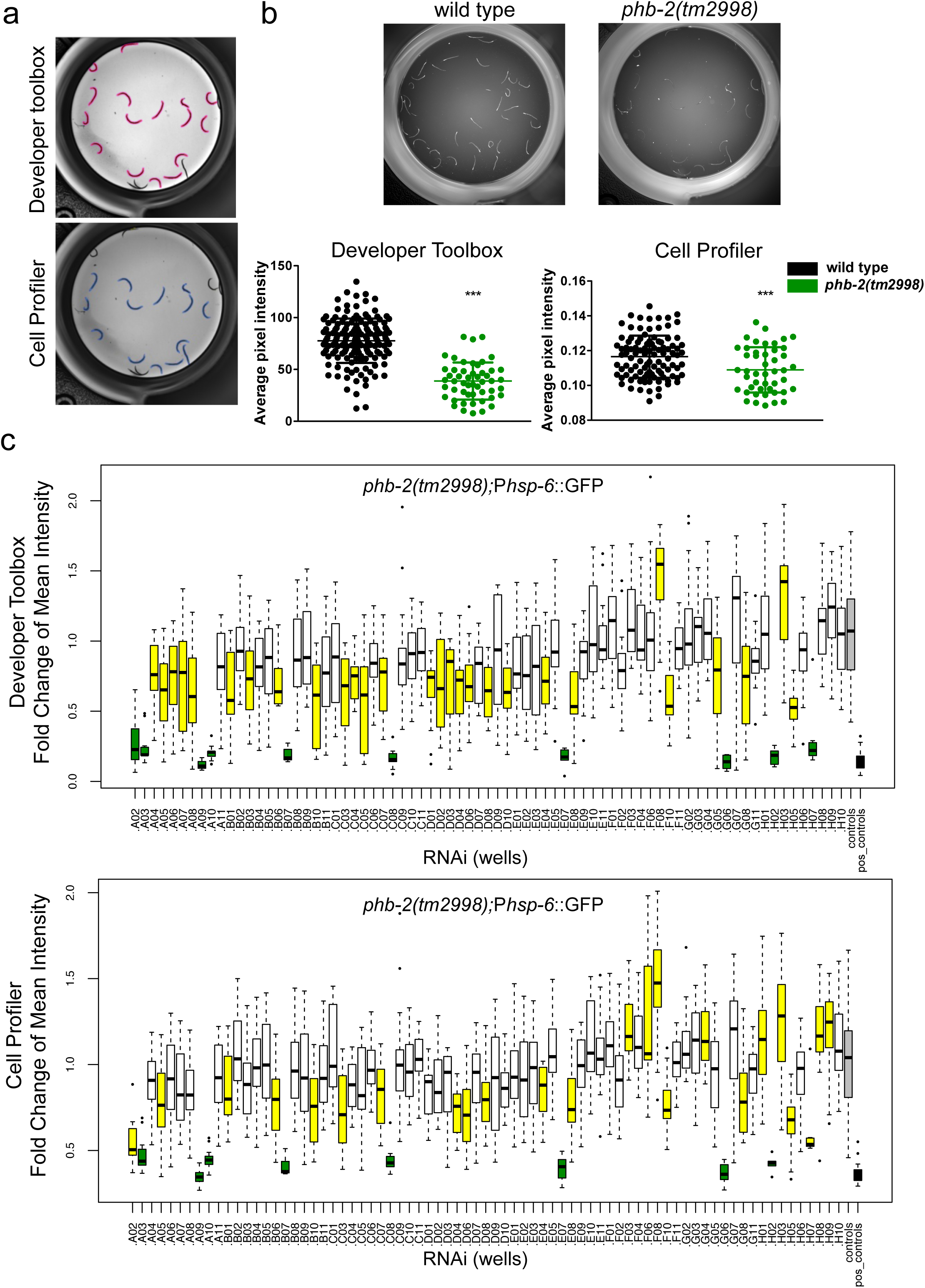
Open sourced segmentation protocol (CellProfiler). **a.** Comparison of the image segmentation output for *phb-2(tm2998)* mutants generated from Developer toolbox (GE Heathcare) versus CellProfiler. **b.** Nile red staining of *phb-2(tm2998)* mutants versus wild type animals. Representative images taken with the IN Cell Analyzer and graphical representation of data coming from the different segmentation protocols. As previously shown [47] depletion of *phb-2* reduces Nile Red staining using both of the protocols. (*** P value < 0.0001; Unpaired t-Test) **c.** The box plots represent fold change mean intensities of the RNAi clones from a randomly chosen 96-well plate from the UPR^mt^ RNAi screen. RNAi clones with a P value < 0.001 appear in yellow and candidates with a FC < 0.5 and a P value < 0.001 appear in dark green. Control wells (control *(RNAi)*) appear in grey and positive control (*atfs-1 (RNAi)*) in black. By comparing both box plots we see that results from the different segmentation protocol are highly similar.

The image analysis protocol generated through the CellProfiler software is provided as Additional file 9, and is easily adaptable to the user’s needs in terms of different fluorescent markers, image formats and image resolutions.

## DISCUSSION

*C. elegans* is an excellent multicellular model for genome wide studies. Its short and well-defined life cycle, its inexpensive and easy maintenance in laboratories and its completely sequenced genome, sharing a high degree of sequence conservation with humans, make *C. elegans* a suitable platform for high throughput and high content screens. Despite all the advantages, large scale studies in *C. elegans* involving essential genes are still not well developed. Temperature-sensitive alleles allow the temporal suppression of gene function, however very few essential genes count with a ts allele [5, 6]. Another possibility is the use of chromosome balancers, which cover already 85% of the genome and a big effort is being carried out for having balancers for all *C. elegans* essential genes by CRISPR [12] and will all be fluorescently labelled (*Caenorhabditis* Genetics Center (CGC), Personal Communication).

Here we present a successful strategy for automated whole animal image-based RNAi screening that can be applied to essential genes when carrying a fluorescently labelled balancer and a gene expression reporter. Developing this automated pipeline is demanding due to two major steps in the workflow. First, sorting homozygous mutant animals expressing a GFP-based stress reporter from a GFP-marked balanced population at an early larval stage (L2) and secondly, a robust automated imaging and segmentation protocol using a microscopy platform not previously utilised for *C. elegans*. Most gene expression reporters in *C. elegans* consist of a given promoter fused to GFP (e.g. the *C. elegans* promoterome [64] or numerous metabolic and stress reporters [44, 65]). By exploiting the Profiler II software feature of the worm sorter, we successfully optimise a protocol for sorting homozygous L2 larvae with an efficiency greater than 95%. We first attempted to sort L1 larvae in order to expose the animals to RNAi from the beginning of larval development. However, the pharynx of the animals occupies approximately one third of the body length at the L1 stage, complicating the profiling. Starting the RNAi treatment at the L2 stage was sufficient to uncover relevant genetic interactions. Nonetheless, interactions that take place during the L1 stage might have been overlooked. COPAS is a convenient approach for high throughput analysis involving balanced strains as it facilitates a task that manually would be impracticable. For balanced strains in the absence of a gene reporter or when the reporter carries a different fluorophore, the sorting of worms does not require the use of the Profiler.

High throughput imaging strategies specific to *C. elegans* have been previously developed using various microscopy platforms [20, 21, 23, 66]. Like other microscopy platforms, the IN Cell Analyzer (GE Healthcare) combines brightfield- and fluorescent-based imaging. We optimize image acquisition to include an entire well of a 96-well plate in a single image, ensuring even brightfield illumination by sealing the 96-well plate with a transparent seal. The optimized autofocus and scanning times of the IN Cell Analyzer ensures image acquisition of an entire 96-well in 2 channels in less than 10 minutes and in 3 channels under 15 minutes. We present here the first automated segmentation protocol for balanced strains in *C. elegans*. We build a user-defined, user-friendly protocol using the Developer Toolbox (GE Healthcare) software. Our segmentation protocol allows intensity-based measurements apart from other measures defined by the user like area, length, curvature, etc., and can be used for any strain without the need of transgenesis as segmentation is done in the brightfield image. One novelty of this segmentation protocol is the measurement of the background surrounding each target. By dilating the targeted worm, we can easily measure and subtract the intensity of the immediate background. This is very appropriate in the cases of dyes that stain plastic and can cause different background intensities depending on the area of the well. Another strong point of our image analysis protocol is the successful identification of worms carrying a pharyngeal GFP element. As COPAS sorting is not 100% efficient, heterozygous worms will develop into fertile adults and lay progeny in the wells. Hence, the need of distinguishing worms carrying the GFP-balancer from homozygous mutants.

The COPAS can serve as an alternative to microscopy-based measurements as it can measure length, optical density and fluorescence emission of single worms [18, 67]. However, image based microscopy platforms have several advantages: they are faster and images are stored, making it accessible for re-analysis. Also, much smaller number of animals can be used for image based assays as compared to the COPAS and thus, allows for exploration of a large number of different treatments in parallel, such as RNAi or drug screens. Another advantage of image-based screens is that multiple outputs can be examined from the image; fluorescent intensities in different channels, size and shape measurements. This helps us to define developmental phenotypes and could be used as well to detect other phenotypes such as sterility by classifying as progeny the small worms (worm length measurement), or to distinguish between thin and fat worms (worm width measurement). We translate the image analysis protocol for CellProfiler [30], a free and open source image-analysis software, making the protocol available to the scientific community. In doing so, we broaden the features of the CellProfiler “WormToolbox” for high-throughput screening of image-based *C. elegans* phenotypes [23, 31]. We provide validation that both protocols produce comparable results.

Prior to the realization of the screen, different optimization steps were needed. Different numbers of worms per well and amounts of bacteria were tested. The final solution is a compromise between having enough worms per well without much overlap, as worms that cross each other are discarded from the analysis. The amount of bacteria needs to be sufficient to avoid starvation, but not too much to prevent anoxic conditions. The speed of shaking during the RNAi treatment has been optimised to prevent the formation of a thick layer of bacteria. Finally, it is worth highlighting the convenience of performing the screen using the same conditions (same batch of plates, same incubator, etc.) to reduce variability.

PHB genes are highly conserved from yeast to mammals. Therefore, to test our automated screening strategy, we perform a chromosome I RNAi screen utilising the OrthoList RNAi library looking for PHB genetic interactors and regulators of the UPR^mt^. The OrthoList RNAi library created by us will be of considerable advantage for *C. elegans* researchers to streamline RNAi screens by focusing on genes with translational potential to human health and reducing screening efforts by 60%.

In *C. elegans*, homozygous *phb-*1 and *phb-2* deletion mutants are sterile and need to be maintained as balanced heterozygous. Here we show that homozygous *phb-2(tm2998)* mutants have delayed development, shorter lifespan and a strong induction of the mitochondrial stress response. Studies looking for the molecular mechanism of this response have identified a number of proteins as essential players for the induction of the UPR^mt^. An uncharacterized temperature-sensitive mutant isolated from EMS, *zc32*, that induces the UPR^mt^ under non-permissive temperature (25ºC) was used to screen chromosome I [44]. RNAi depletion of two genes, *lin-35* and *ubl-5* suppressed induction of P*hsp-6*::GFP and P*hsp-60*::GFP. With the same mutant, three other components of the UPR^mt^ were described, the transcription factors DVE-1 and ATFS-1 and the mitochondrial transporter HAF-1 [45, 49]. HAF-1 was at first suggested as an essential upstream component of the UPR^mt^ [49, 50], however conditions that induce the UPR^mt^, such as inhibition of mitochondrial protein import [68] or RNAi against the cytochrome *c* oxidase subunit *cco-1* do not require HAF-1 [50, 52]. We here demonstrate that, rather than blocking, loss of HAF-1 further induces *hsp-6* expression upon depletion of *phb-1*, *phb-2* or *spg-7* (Figure 1d and Additional file 2). These data suggest that more studies are required for the complete understanding of the regulation of the mitochondrial stress response. More recently, Shore *et al.* analysed the induction of different cytoprotective responses such as ER stress, mitochondrial stress, oxidative and osmotic stress, among others, upon depletion of 160 genes reported to increase lifespan in *C. elegans* [62]. They identified 42 RNAi clones required for the activation of the UPR^mt^ triggered by antimycin, a chemical that disrupts complex III of the mitochondrial electron transport chain (ETC). In a genome wide screen, Runkel *et al.* found 55 genes necessary for the activation of the UPR^mt^ induced by paraquat treatment [61]. Applying our protocol, we describe 208 genes that when depleted reduce the mitochondrial stress response in *phb-2* mutants. On comparison with the previously published data, we found 45.5% of genes previously identified to be required for the activation of the UPR^mt^. This big number of additional candidates can be explained by the quantitative nature of the protocol. In addition, genes affecting development might result in reduced UPR^mt^ because P*hsp-6*::GFP expression increases as worms develop. Similarly, it is worth stressing the different experimental conditions: chemical treatments or a temperature-sensitive mutant versus a mutant of the mitochondrial PHB complex, which might trigger the mitochondrial stress response through an alternative pathway. In addition to earlier published clones, we identify numerous genes encoding proteins of the two ribosomal subunits, proteins associated with protein transport including nuclear importins, proteasomal subunits and genes involved in mRNA processing, all processes already known to regulate the UPR^mt^.

Interestingly, we encounter 303 PHB interactors whose depletion causes a developmental defect. The fact that Chromosome I (left) is particularly enriched in essential genes partially explains this high number of interactions [69]. We uncover 31% of the genes previously described to affect development [57, 63]. As mentioned before, experimental procedures are different and could explain the non-complete replicability: *phb-2* mutants have *per se* a developmental delay and RNAi treatment is performed from L2s instead of from eggs. Therefore, L1 acting genes could have been missed. Interestingly, we find several genes involved in fatty acid metabolism, which is mostly carried out in the mitochondria, genes encoding for nuclear pore complex proteins as well as genes implicated in DNA repair. It would be of a very high impact to study more in detail why genes involved in these processes affect the development of prohibitin mutants.

## CONCLUSIONS

The function of many essential genes is not well understood. The method described here combines automated worm sorting and high-content image analysis that can be adapted and applied to any balanced strain carrying a fluorescently labelled balancer and carrying or not an additional gene expression reporter. Therefore, this method can be instrumental to increase our knowledge on the biology of essential genes and the generation of genetic interaction networks involving essential genes in *C. elegans*. Approximately 60% of *C. elegans* essential genes have human orthologues, making the worm relevant to study the function of essential genes that will impact human health.

## METHODS

A detailed description of the protocol, day by day, is Additional file 3.

### *C. elegans* strains and maintenance

The *C. elegans* strains used in this study are: N2 (wild type), MRS106 *phb-2(tm2998)*/*mIn1*[*dpy-10(e128)mIs*14(P*myo-2*::GFP)]II;*zcIs13*[P*hsp-6::*GFP]V, MRS104 *phb-2(tm2998)*/*mIn1*[*dpy-10(e128)mIs*14(P*myo-2*::GFP)]II;z*cIs13*[P*hsp-60::*GFP]V, MRS50 *phb-2(tm2998)*/*mIn1*[*dpy-10(e128)mIs*14(P*myo-2*::GFP)]II;*haf-1(ok705)*IV;*zcIs13*[P*hsp-6::*GFP]V, BR6118 *haf-1(ok705)*IV;*zcIs13*[P*hsp-6::*GFP]V, BR6115 *phb-1(tm2571)I/hT2[bli-4(e937) qIs48*(P*myo-2*::GFP)*](I;III)*, 10 times outcrossed before introducing the *hT2* balancer, BR6108 *phb-2(tm2998)/mln1*[*dpy-10(e128) mIs*14(P*myo-2*::GFP)]*II*, 10 times outcrossed before introducing the *mIn1* balancer, PE255 *feIs5[sur-5::luc+::*GFP*;rol-6(su1006)]V,* MRS229 *phb-2(tm2998)/mln1*[*dpy-10(e128)mIs*14(P*myo-2*::GFP)]*II;feIs5[sur-5::luc+::GFP;rol-6( su1006)]X.*

Unless otherwise stated, we culture the worms according to standard methods [70]. We maintain nematodes at 20°C on NGM agar plates seeded with live *Escherichia coli* OP50 (obtained from the *Caenorhabditis* Genetics Center (CGC)). To obtain synchronized L1 larvae, we collect eggs by hypochlorite treatment and allow them to hatch and arrest by overnight incubation in M9 at 20 ºC with agitation.

### Generation of the OrthoList RNAi sub-library

Plates from Ahringer’s library containing the clones of interest (purchased from MRC Geneservice) are replicated to LB agar supplemented with ampicillin (100 μg/ml - Sigma-Aldrich) and tetracycline (15 μg/ml – Sigma-Aldrich) using the pin replicator (BOEKEL) and grown overnight at 37ºC. The day after, the selected clones are inoculated in 1.3 ml of LB supplemented with ampicillin (100 μg/ml - Sigma-Aldrich), tetracycline (15 μg/ml – Sigma-Aldrich) and glycerol 8% in deep well plates (VWR) and incubated overnight at 37ºC with shaking (180 rpm - New Brunswick™ Innova^®^ 44/44R). Last column of the plates is left free for the convenient controls. Next day, 120 μl of the culture is transferred to microtiter plates using the Precision XS Microplate Sample Processor (BIOTEK) and frozen at −80ºC.

### Preparation of the bacteria

Using a pin replicator (BOEKEL), we replicate the plate in LB agar supplemented with ampicillin (100 μg/ml - Sigma-Aldrich) and tetracycline (15 μg/ml – Sigma-Aldrich). Bacteria are grown overnight at 37ºC. The advantage of growing the bacteria in solid media is the ease to visualize the clones where the bacteria do not grow. If needed, it can be kept at 4ºC for 2 days maximum. Next day, the RNAi library is inoculated in 2.2 ml - 96 well plates (VWR). Using the pin replicator we inoculate the bacteria from LB agar into 1.2 ml of LB supplemented with ampicillin (100 μg/ml - Sigma-Aldrich) and tetracycline (15 μg/ml – Sigma-Aldrich). Positive and negative controls are added in the last column of the plate. The bacteria are grown overnight at 37ºC with shaking (180 rpm - New Brunswick™ Innova^®^ 44/44R). In order to have fresh cultures, the day of the sorting, 100 μl of the O/N cultures are inoculated in 900 μl of LB supplemented with ampicillin and tetracycline in deep well plates (VWR) and incubated for 3 hours at 37ºC with shaking. IPTG (1 mM– Sigma-Aldrich) is added to the wells to induce the expression of the plasmid for 2 hours at 37ºC with shaking. The cultures are harvested by centrifugation (10 minutes, 3200 g, 4ºC – Eppendorf 5810R) and pellets are resuspended in 250 μl of S-medium supplemented with carbenicillin (25 μg/ml – Sigma-Aldrich), IPTG (1 mM – Sigma-Aldrich) and cholesterol (5 μg/ml – Sigma-Aldrich).

### Worm preparation and sorting

For each round of sorting, worms are synchronized obtaining eggs from gravid hermaphrodites by hypochlorite treatment. Briefly, 20 ml of liquid culture with worms suspended in S-medium supplemented with OP50 (30 g/l wet weight) are washed with M9 until the supernatant appears clear of bacteria. Bleaching solution is added and tubes are energetically agitated for 2 minutes. After centrifugation and removal of the supernatant, the worms are washed with M9. A second round of bleaching solution is added for less than 1 minute. The pellet is then washed three more times with M9 and filtered with 40 μm Nylon Cell Strainers (VWR) to remove the possible remains of adult worms. Embryos are allowed to hatch overnight in M9 at 20ºC with shaking (120 rpm - New Brunswick™ Innova^®^ 44/44R). The following day, starved L1s are placed in S-medium with OP50 (30 g/l) for 48 hours at 20ºC with shaking (120 rpm - New Brunswick™ Innova^®^ 44/44R). At that point, the population is heterogeneous, since the homozygous *phb-2(tm2998)* mutants show a developmental delay, and are at the second larval stage. Worms are washed out from the OP50 culture by successive centrifugations until the supernatant is clear. Worms are resuspended in M9 supplemented with 0.01% Triton X-100 (T8787, Sigma-Aldrich) to avoid sticking to the plastic. Next, we sort 40 worms per well using enhanced mode, with a sort delay (time from analysis of the object to the sort command) of 7 milliseconds and a sort width (drop volume) of 6 milliseconds.

Subsequently, 25 μl of S medium supplemented with carbenicillin (25 μg/ml – Sigma-Aldrich), IPTG (1 mM – Sigma-Aldrich) and cholesterol (5 μg/ml – Sigma-Aldrich) are added to the worms. Finally, 75 μl of the bacterial culture is added. Worms are incubated for 48 hours at 20ºC with shaking (120 rpm - New Brunswick™ Innova^®^ 44/44R) until they reach the young adult stage.

We aim to keep the sheath flow rate constant at 9.5 ml/min and a worm concentration of 15-20 events per second. At the start of each experiment, a small sample (1 single worm in 96 wells) is sorted and visually verified to confirm a correct sorting, that is, correct number of animals and correct selection of the population.

The worm sorter is a pressurized machine and one should pay attention to any clog that might interfere with the liquid flow from the sample cup to the flow cell.

Every day, before starting, the tubes are cleaned by passing consecutively from the sample cup 10% bleach, water and 70% ethanol, and then rinsed out with abundant water. Moreover, all the solutions are passed through a 40 μm Nylon Cell Strainers (VWR) filter.

For imaging in the red channel, homozygous *phb-2(tm2998)* mutants are sorted at the L1 larval stage after overnight starvation, using the COPAS Biosort without utilizing the Profiler feature, into 96-well plates. Wild type animals are pipetted manually. Approximately 40 worms are grown in 100 *µ*l HT115 (DE3) bacteria containing the empty vector pL4440 supplemented with 100 nM Nile Red and incubated at 20°C with shaking.

### Imaging of multiwell plates

In order to have clear images, the plates are washed by sequential flush of water, shaking to disaggregate the bacteria, sedimentation of the worms and aspiration of the supernatant (EL406 washer dispenser, BioTek). Prior to this, 10 μl of Tetramisole hydrochloride (100 mM - Sigma-Aldrich) are added to each well to paralyze the worms. Each well is filled to brim and sealed with transparent sealplate (SIGMA), to ensure horizontal meniscus required to give uniform brightfield illumination across each well. Pictures in brightfield, green and/or red channel are acquired with the IN Cell Analyzer 2000 (GE Healthcare) using a 2x objective in order to have the entire well in one image. A whole 96 well plate can be imaged in two channels in less than 10 minutes and in three channels in less than 15 minutes.

### Data analysis

Our pipeline consists of several filtering steps, quality control and statistical test. First, based on the green head ID, worms with a green pharynx, are discarded, as well as worms with a length smaller than 550 μm. In order to remove outliers, the 5^th^ and the 95^th^ percentile of the distribution are excluded. After filtering, wells with less than 5 worms are removed from the analysis. Bacteria containing an empty vector, pL4440, is used as negative controls and *atfs-1(RNAi)*, that suppresses almost completely the UPR^mt^ in *phb-2* mutants, as positive control (Figure 5a). A quality assay is performed in the control wells: only control wells with mean GFP intensity between 200 and 500 arbitrary units (a.u.) and a coefficient of variation < 0.5 are accepted. If less than 2 control wells remain accepted, the plate is discarded and repeated. In order to make data from different plates comparable, data are normalized by dividing the GFP value of each worm by the mean of the GFP of the four negative control wells. Finally, statistics are assessed by running an ANOVA test followed by a Dunnett’s test. Candidates are defined based on the P value and the fold change (FC) (P value < 0,001 and FC < 0.66 or FC > 1.5).

Interaction networks are built using String (http://string-db.org), based in predicted and described interactions in different organisms and functional annotation clustering are performed with DAVID Bioinformatic Resource 6.8 (https://david.ncifcrf.gov/).

### Western blot analysis and antibodies

Protein levels are quantified by immunoblot assay. A synchronized population of worms is grown at 20°C until they reached young adult stage. Worms are transferred to NGM plates without food and allowed to crawl for half an hour in order to remove excess of bacteria and 40 animals are collected in 15 *µ*l of M9. The same volume of 2x sample buffer (100 mM Tris pH 6.8, 0.02 % Serva blue G, 8 % SDS, 24 % glycerol and 4 % mercapto-ethanol) is added, sample is boiled for 5 min, spun for 1 min (4°C) and kept at −80°C for 3 - 5 days. 16 *µ*l of sample is run in 12% SDS-PAGE. Following electrophoresis, proteins are transferred to a PVDF membrane (Immobilon-P; Millipore Cat. No. IPVH00010). The immunoblots are visualized by chemiluminescent detection (Pierce ECL Western Blotting Substrate; prod # 32209). Western blots are incubated with APP-2 antibody (1:3000, overnight at 4°C) and anti-actin as described [51].

### Luciferase assay to determine developmental rates

We use the reporter strains PE255 and MRS229 to measure larval developmental timing. In order to ensure all animals start development at the same time, arrested L1s are first manually pipetted to a white 96-well plate, 1 worm per well, containing 100 *µ*l of S-basal with 100 *µ*M D-Luciferin. Development is resumed by addition of 100 *µ*l of S-basal with 20 g/l *E. coli* OP50 and 100 *µ*M D-Luciferin. Plates are sealed with a gas-permeable cover (Breathe Easier, Diversified Biotech). Luminescence is measured in Berthold Centro LB960 XS3 for 1 sec, typically at 5-min intervals. Experiment is done inside temperature-controlled incubators (Panasonic MIR-154). The raw data from the luminometer is analysed as in Olmedo, M. *et al.* [48]. Briefly, the raw data is trend-corrected and thresholded using 75% of the moving average to produce a binarized output in order to determine onset and offset of the molts. Data is evaluated for onset and offset of molting by detecting the transitions in the binarized data.

### Lifespan Analysis

All lifespans are done at 20°C. Synchronized eggs are obtained by hypochlorite treatment of adult hermaphrodites and placed on NGM plates containing OP50 *E. coli* bacteria. During the course of the lifespan, wild type adult nematodes are transferred every day during their reproductive period and afterwards on alternate days. Homozygous *phb-1(tm2571)* and *phb-2(tm2998)* mutants are separated from their respective heterozygous populations at L3 stage to a separated plate and transferred every alternate day. Worms are scored as dead when they stop responding to prodding, while exploded animals, those exhibiting bagging, protruding gonad or dried out on the edge of the plates are censored. GraphPad Prism software is used to plot survival curves and significant differences in lifespan are determined by using the log-rank (Mantel–Cox) test.

### Slide Imaging

A semi-synchronous embryo population is transferred to NGM plates seeded with the appropriate RNAi bacterial clone. Animals are allowed to grow at 20ºC until young adult stage when 20-30 worms are mounted on 2% agarose pads in M9 medium containing 10 mM Tetramisole hydrochloride (Sigma) and imaged using an AxioCam MRm camera on a Zeiss ApoTome Microscope. In order to remove excess of bacteria worms are first transferred to NGM plates without food and allowed to crawl for half an hour. Emission intensity is measured on greyscale images with a pixel depth of 16 bits. Image analysis is performed using the ImageJ software and data are analysed by one-way ANOVA using GraphPad Prism. One independent replicate out of three is shown.

## DECLARATIONS

### Ethics approval and consent to participate

Not applicable.

### Consent for publication

Not applicable.

### Availability of data and material

All data generated or analysed during this study are included in this published article and its additional files. Raw data can be found in Additional files.

Additional files 4 and 8: Worm segmentation protocols for Developer Toolbox

Additional files 5 and 6: Raw values for the RNAi screen of Chromosome I (P*hsp-6*::GFP and worm size) corresponding to Figure 5.

### Authors’ contributions

MAS conceived and design the study. MAS, BHR, APE, MJRP, BS and MO performed experiments. MAS backcrossed and balanced strains. BS performed western blots and crossings (guided by MAS) at RB laboratory. MO, APE and MJRP measured developmental phenotypes. MAS, BHR, APE, and MJRP optimised sorting protocols. VM developed segmentation protocols using Developer Toolbox (GE, Healthcare), guided by BHR, APE, MJRP and MAS. BHR and APE performed the experiments and analysed the data (guided by MAS). PA, MM, MAS, MJRP and SGH designed and generated the OrthoList RNAi library. BHR optimised the CellProfiler protocol. BHR, APE and MAS wrote the manuscript with input from VM. All authors read, commented and approved the final manuscript.

## Acknowledgements

We thank Carolina Wählby for her invaluable support translating the image analysis protocol to CellProfiler and for enlightening comments on the manuscript and Juan Tena for writing the pipeline in R for computerising the analysis of the UPR^mt^ screen. We acknowledge both, the *Caenorhabditis* Genetics Center (CGC) and the *C. elegans* National Bioresource Project of Japan (NBRP) for providing strains used in this study. In particular, prohibitin deletion alleles were provided by the NBRP. Special thanks to Liesbeth de Jong for illustrating the screening strategy (Figure 2).

---

**Additional File 1, Figure S1: Effect of *haf-1* deletion in the UPR^mt^.**

**Additional File 2, Table S1: Day by day description of the experimental procedure.**

**Additional File 3: Detailed description of the protocol, day by day, together with tips and discussion of the optimization of the approach.**

**Additional File 4: Break down of the image analysis protocol for green images.**

**Additional File 5: Raw data of P*hsp-6*::GFP reporter measurements.**

**Additional File 6: Raw data of size measurements.**

**Additional File 7, Figure S2: Outline of the combined green / red image analysis for balanced mutants.**

**Additional file 8: Break down of image analysis protocol for green and red measurements.**

**Additional file 9: Break down of CellProfiler protocol for green and red measurements.**

